# Integrative meta-analysis reveals that most yeast proteins are very stable

**DOI:** 10.1101/165290

**Authors:** Keira Wiechecki, Sandhya Manohar, Gustavo Silva, Konstantine Tchourine, Samson Jacob, Angelo Valleriani, Christine Vogel

## Abstract

Measurements for protein half-lives in yeast *Saccharomyces cerevisiae* reported large discrepancies, with median values between minutes to several hours. We present a unifying analysis that provides a consistent half-life estimate, based on our re-analysis of three published and one new dataset of cells grown under similar conditions. We found that degradation of many proteins can be approximated by exponential decay. Protein disappearance was primarily driven by dilution due to cell division, with cell doubling times ranging from ∼2 to 3.5 hours across the four experiments. After adjusting for doubling time, protein half-lives increased to median values between ∼7.5 to ∼40 hours. Half-lives correlated with cell doubling time even after adjustment, implying that slow growth also slows protein degradation. All estimates were validated by multiple means and were robust to different analysis methods. Overall, protein stability correlated with abundance and showed weak enrichment for degradation signals such as degrons and disordered regions. Long-lived proteins often functioned in oxidation-reduction and amino acid synthesis. Short-lived proteins often functioned in ribosome biogenesis. Despite some overall differences in behavior, all methods were able to resolve subtle difference in half-lives of ribosomal proteins, e.g. the short lifespan of RPL10. Finally, our results help the design of future experiments: time series measurements need to cover at least two to three cell doubling times for accurate estimates, exponential decay provides a reasonable proxy for protein stability, and it can be sufficiently estimated with four measurement points.

## Introduction

Protein degradation is, next to transcription, translation, and RNA stability, a major route to fine-regulate the abundance of proteins. Ubiquitination and subsequent proteasomal degradation are the major pathway regulating protein breakdown (Lee and Goldberg, 1998), in addition to lysosomal degradation. However, attempts to identify sequence motifs that indicate ubiquitination have so far failed (Kim et al., 2011) and other degrons, such as the PEST motif or the N-terminal amino acid also have very small predictive power of protein half-life (Bachmair et al., 1986; Gsponer et al., 2008; Kristensen et al., 2013; Rechsteiner and Rogers, 1996).

Therefore, experimental approaches that measure protein stability, e.g. in form of protein half-life, are essential to further our understanding of this mode of regulation. For the budding yeast *Saccharomyces cerevisiae*, several large-scale studies have examined protein half-lives of wild-types strains growing in log-phase and rich medium. Earlier work used cycloheximide and western blotting and identified a median half-life of ∼45min across >3,000 proteins (Belle et al., 2006). Later studies used less invasive approaches and monitored the disappearance of isotopically labeled proteins by mass spectrometry (Ong et al., 2002).

Surprisingly, while these studies used almost identical methods (**Suppl. Table S2**), they provided vastly different half-life estimates. For example, Christiano et al. monitored protein stabilities over three hours (180 min) and found median half-lives of approximately nine hours (Christiano et al., 2014). In contrast, Martin-Perez et al. conducted a six-hour experiment in both auxotroph and prototroph yeast and found protein half-lives close to the cell doubling time of ∼two hours (108 min)(Martin-Perez and Villen, 2015).

To resolve these discrepancies, we set out to analyze the data with identical methods to derive a consistent estimate of protein half-lives in yeast and to maximize cross-study comparability. To do so, we processed the Christiano, Martin-Perez, and our own data in the same way. We identified challenges in cross-comparison of the dataset due to the impact of cell doubling time, sparse sampling over short time periods, and the inherent measurement noise. However, we were able to resolve the different findings and show that the four datasets -Christiano, Martin-Perez Prototroph and AuxoTroph, Silva -agree in major trends and characteristics, such as a positive correlation of stability with protein abundance and function biases. Dilution due to cell division explains most of protein disappearance from the cell. Once one corrects for cell doubling time, protein half-lives average to many hours. Therefore, future experiments need to monitor the proteome for at least two to three cell doublings.

## Results

### Many proteins can be well-modeled by first-order decay

To derive consistent protein half-life estimates, we analyzed four datasets describing a total of >3,300 proteins from wild-type yeast *Saccharomyces cerevisiae* grown under similar conditions in rich medium (**Suppl. Data 1**). All studies used pulsed SILAC proteomics to measure proteins with different isotopic labels over the course of the experiment. To increase comparability even further, we selected four time points from each time series measurement that covered a similar range of values, i.e. approximately 4 hours (240 min, **Table 1**). We named the datasets after the first author of the study, i.e. Christiano (Christiano et al., 2014), Martin-Perez (Martin-Perez and Villen, 2015), and Silva (this study), respectively. Remaining differences are listed in **Suppl. Table S2** and are discussed in a later section. The Martin-Perez datasets are present in duplicate and contain data for up to 8 hours (480 min) which we also discuss below.

**Table 1:** Most proteins are well-modeled by exponential degradation. Protein half-life varies across genes and across experiments. Unadjusted *T*_1/2_* is the half-life calculated by the exponential model prior to adjusting for cell division time *T*_*CC*_. Adjusted *T*_1/2_ is the half-life calculated by the model after accounting for *T*_*CC*_ (see Methods). Proteins for which *T*_1/2_* < *T*_*CC*_ were considered outside the appropriate measurement range, as adjusting these half-lives for doubling time results in nonsensical values.

|  | Christiano | Silva | Martin-Perez<br>Prototroph | Martin-Perez<br>Auxotroph |
| --- | --- | --- | --- | --- |
| **Doubling time $T_{CC}$ (min)** | 150 | 208 | 122 | 108 |
| **Measurement time points (min)** | 30, 60, 120,<br>180 | 30, 90, 150,<br>240 | 30, 60, 90, 240 | 30, 60, 90,<br>240 |
| **Total number of proteins measured** | 3,518 | 2,214 | 1,029 | 1,194 |
| **Number of proteins modeled by first-order (exponential) decay (LOOCV<0.01 and R2>0.9)** | 3,067<br>(87%) | 1,374<br>(62%) | 475<br>(46%) | 579<br>(48%) |
| **Median $T_{1/2}^*$ of exponentially degrading proteins (min)** | 120 | 256 | 112 | 104 |
| **Range $T_{1/2}^*$ (min)** | 42 - 561 | 118 - 449 | 61 – 1,938 | 56 – 2,551 |
| **Median $T_{1/2}$ of exponentially degrading proteins, adjusted for cell doubling (min)** | 609 | 2,357 | 454 | 657 |
| **Number of proteins modeled by first-order decay, but with half-lives outside measurement range** | 24<br>(0.01%) | 1,124<br>(51%) | 150<br>(15%) | 192<br>(16%) |

To ensure comparability between datasets and account for variability within the time series measurements, we processed all data identically and applied the same cutoffs. Briefly, we scaled the data using ratios of abundance of newly synthesized protein to abundance of proteins synthesized prior to the start of the time series (**Methods**). All results are robust even if we scaled based on highly abundant and stable proteins or no scaling was used (**Suppl. Notes and Figure S1**).

To further increase consistency in our comparison of the four datasets, we focus on proteins whose degradation followed a first-order decay function, implementing exactly the same method described in Schwanhaeusser and Christiano (Christiano et al., 2014; Schwanhausser et al., 2011), using the same cutoffs. As discussed below and in recent literature, protein degradation likely follows a function more complex than first-order decay (McShane et al., 2016). However, such functions can only be monitored by dense temporal sampling with more than four or even six or eight time points as used in the present studies. Therefore, for the purpose of this analysis, i.e. to consolidate the very different half-life estimates across different studies, we focus on the subset of proteins whose degradation is well-approximated by exponential decay, i.e. ∼60 to 90% of the proteins in the datasets (**Table 1**). Fewer proteins followed exponential decay when we used more time points, confirming that the ‘real’ decay function is not exponential: only 35% and 24% of proteins were well-modeled in the extended Martin-Perez dataset with six or eight time points, respectively (**Table 1**). Exponential decay offers a reasonable approximation of protein stability useful for many purposes, but it does not appear to be the primary underlying function.

### Yeast proteins are very stable

We calculated two half-lives for each protein, *T*_1/2_* and *T*_1/2_, which were either not adjusted or adjusted for the contribution of cell division to the disappearance of the protein in the cell, using identical methods. All calculations and datasets produced a wide range of values, from ∼45min to several hours (**Figure 1**). Therefore, we first compared median values from each datasets. Median *T*_1/2_*, which is not adjusted for doubling, ranged from 2 to 4 hours (120 to 256 min, **Figure 1, Table 1**). Indeed, the values for the individual datasets were very close to the cellular doubling time in the respective datasets which ranged from 122 to 208 min. For example, cells used by Silva divided much less frequently than the other yeast strains and the proteins lived much longer than proteins in the Martin-Perez or Christiano datasets. These results indicate that many proteins were very stable and mostly disappeared from the cell due to cell division. Indeed, once we adjusted for the contribution of cell doubling (*T*_*CC*_), the median half-lives were much longer, ranging from 7.5 to 40 hours (**Table 1**). Intriguingly, half-lives still correlated with doubling times even after adjusting for cell division, suggesting that the cell’s growth rate affects stability in addition to the effects of dilution (**Figure 1**). In other words, in the slow-growing Silva strain, proteins were particularly resistant to proteasomal degradation compared to other strains. The relationship held true even for shorter measurement ranges (**Suppl. Figure S2**).

**Figure 1:**
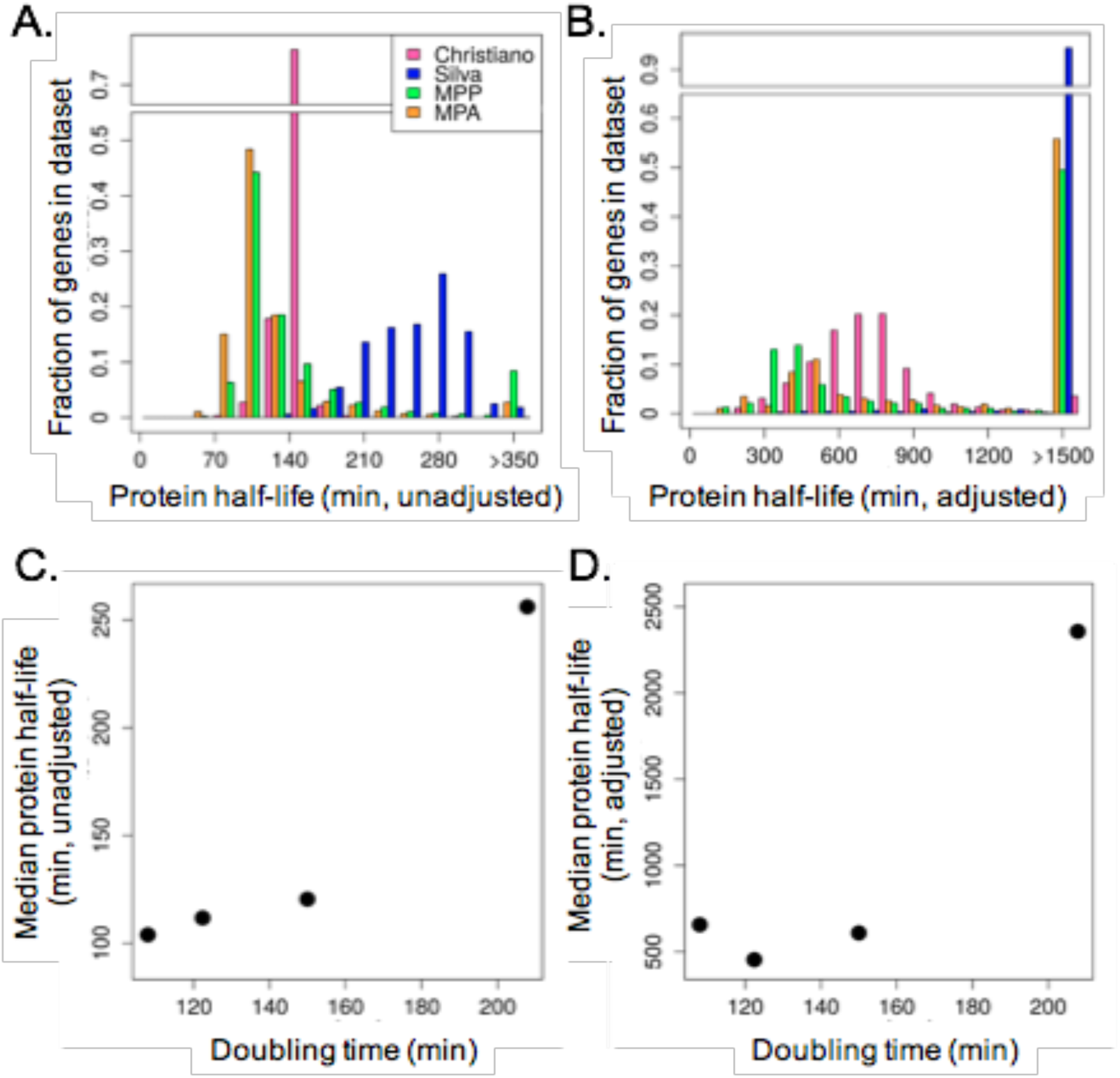
Protein stability correlates with yeast cell division time. **A., B.** Frequency distribution of protein half-lives not adjusted and adjusted for doubling time (*T*_1/2_* and *T*_1/2_, respectively). The rightmost column includes all proteins with half-life greater than the maximum value shown on the graph. Extended results are shown in **Suppl. Figures S1 and S2. C., D.** The doubling time *T*_*CC*_ for the three datasets is plotted against the median protein half-life which is unadjusted (*T*_1/2_*) or adjusted (*T*_1/2_) for cell division time. MPP – Martin-Perez Prototroph; MPA – Martin-Perez Auxotroph

These results highlight the biggest challenge in studying protein half-lives: since many proteins are very stable, it takes several hours for a measurable fraction to be degraded. If the measurement ends too soon, degradation of long-lived proteins cannot be assessed. Even measurements over 3 to 4 hours (i.e. 180 to 240 min as performed by Christiano and Silva), were too short: stable proteins would not ‘disappear’ enough during this time to be reliably measured, and therefore often result in ‘infinite’ half-lives. These infinite half-lifes do not mean that the proteins are immortal, but that the experimental design could not assess their stability. Indeed, up to 50% of the proteins degraded so slowly that their half-life could not be estimated (**Table 1**).

For these reasons, we also calculated half-lives in the only dataset that covered up to 6 hours (360 min, Martin-Perez). Note that Martin-Perez et al. also collected an 8 hour (480 min) time point, but that did not deliver a large number of proteins following exponential decay. A substantial fraction of the proteins in the 360 min time course was well-modeled by exponential decay, and properties of these proteins were consistent with those from the main estimates covering 240 min time courses (**Suppl**. **Table S1, Figure S9**)-increasing our confidence in the generalizability of our results.

However, comparing half-life estimates for the same protein in the same study using the 240 versus a 360 min time series data showed another challenge: the 240 min time course overestimated half-lives for long-lived proteins, i.e. those with half-lives longer than ∼500 min (**Suppl. Figure S9**). Therefore, we suggest that pulsed SILAC time series experiments should cover a minimum of two and ideally at least three cell doubling times to accurately measure degradation of long-lived proteins.

### Common trends emerge

The rigorous consistency and filtering resulted in protein half-life estimates that show consistent characteristics, suggesting that the estimates might reflect true values. First, while different at an absolute scale, half-life estimates correlate between datasets (**Figure 2**). While the trend is most visible for binned data, it is statistically significant also when individual values are compared (p-value < 0.01).

**Figure 2:**
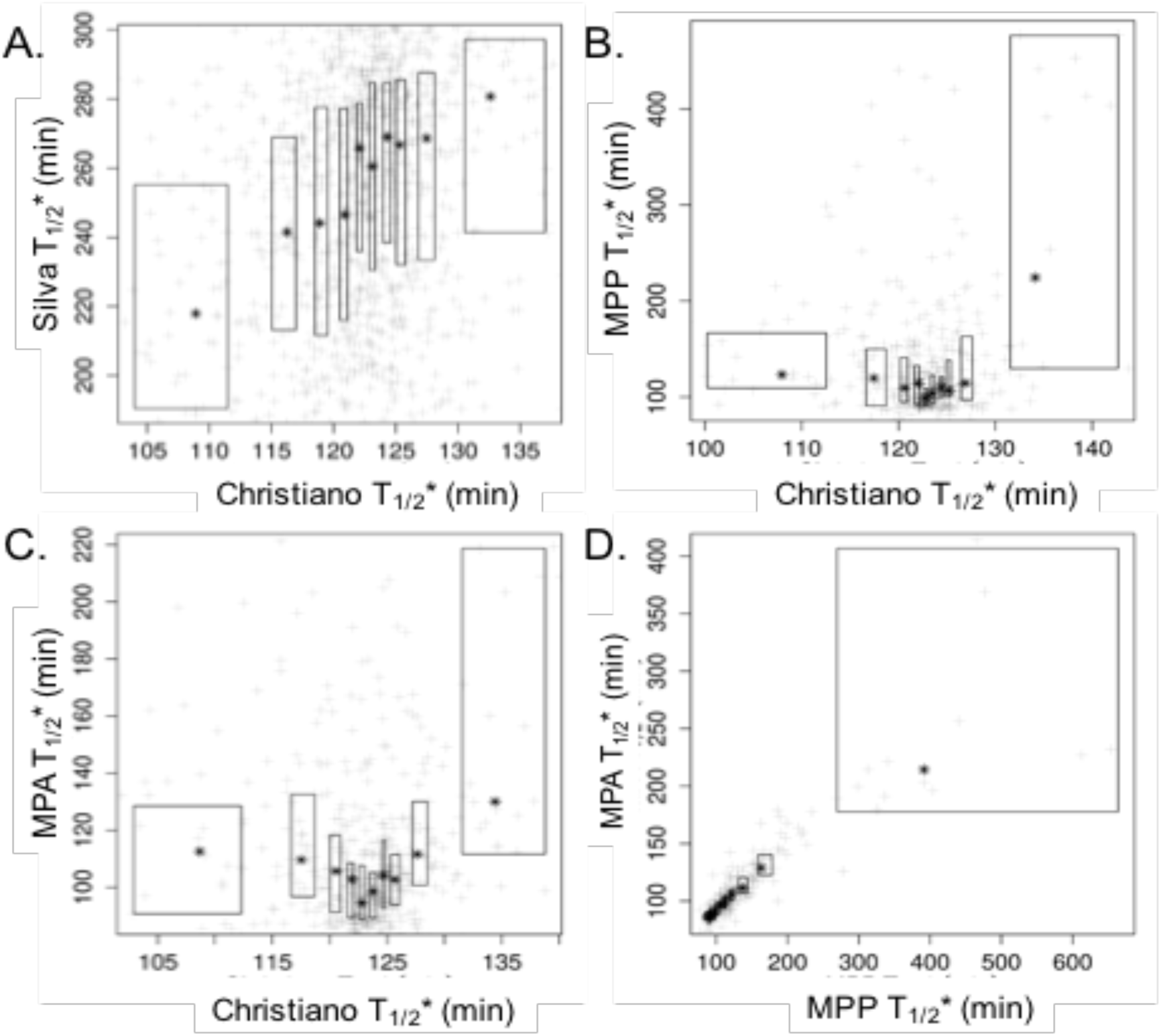
The datasets agree with respect to general trends. The scatter plots show the the relationship between unadjusted protein half-lives (*T*_1/2_*) from the four datasets. Data were rank-ordered and then split into ten equal size bins. Black stars indicate the median values in each bin. Boxes indicate the second and third quartiles within the bins; stars indicate the median values. The total range of the data are shown in **Suppl. Figure S4.** Correlations were calculated for all data points representing Pearson’s r and Spearman’s ρ. **A.** Christiano *T*_1/2_* vs. Silva *T*_1/2_*; r = 0.33 (p < 0.01) and ρ = 0.29 (p < 0.01). **B.** Christiano *T*_1/2_* vs. Martin-Perez prototrophic *T*_1/2_*; r = 0.49 (p < 0.01) and ρ = 0.12 (p = 0.01). **C.** Christiano *T*_1/2_* vs. Martin-Perez autotrophic *T*_1/2_*; r = 0.68 (p < 0.01) and ρ = 0.12 (p < 0.01). **D.** The Martin-Perez Prototroph (MPP) and Martin-Perez Auxotroph (MPA) data strongly correlate, indicating that prototrophs and auxotrophs are similar in protein half-lives. r = 0.70 (p < 0.01) and ρ = 0.90 (p < 0.01).

Second, we observe consistent function enrichments across the datasets (**Table 2**). As seen previously (Martin-Perez and Villen, 2015), long-lived proteins were enriched for the oxidation-reduction, Tri-Carboxylic Acid cycle and amino acid synthesis. Short-lived proteins were enriched for RNA processing and ribosome biogenesis.

**Table 2:** Long- and short-lived proteins share functional enrichment between datasets. The table illustrates enriched functions in 50% longest-and shortest-lived proteins **(A., B.**, respectively) with the average false discovery rate <0.05 in the Christiano and Silva and <0.1 the Martin-Perez datasets (**Suppl. Dataset 2**).

| <u>Significantly enriched function</u> | <u>Example genes</u> |
| --- | --- |
| **<u>A. Long-lived proteins</u>** |  |
| Oxidation / reduction ('De novo' purine biosynthesis, Pyruvate metabolism, GTP biosynthesis) | ADE16 ADH1 ADH6 AIM17 ALD3 ALO1 ARI1 CBR1 CCP1 CIR2 DLD1 DLD3 GLC7 GRE3 HOM6 HYR1 IDP1 IFA38 IMD2 IMD3 IMD4 LYS9 MCR1 PAN5 PDB1 PDX3 PET9 PGA3 PGM2 PRO2 QCR2 RNR2 SFA1 SOD2 TDH1 TDH2 TPA1 YCP4 |
| Amino acid biosynthesis | ALD3 ARO2 ARO3 ARO4 ARO7 ARO8 ASN2 BAT1 BAT2 CAR2 CYS4 FPR1 HOM3 HOM6 IDP1 ILV1 ILV2 LYS21 LYS9 MET17 MRI1 PRO2 SER1 SHM1 THR4 TRP2 TRP3 |
| **<u>B. Short-lived proteins</u>** |  |
| Ribosome biogenesis (rRNA processing, ribosome subunit) | CDC73 CRM1 EMG1 FUN12 GCD11 GLC7 GSP1 MEX67 NSA1 NSP1 PAT1 PUB1 RNA1 RPB5 RPG1 RPL10 RPL25 RPL3 RPL30 RPL5 RPL6A RPL6B RPL7A RPL7B RPL8A RPL8B RPO26 RPP0 RPS13 RPS15 RPS1A RPS1B RPS2 RPS20 RPS21A RPS3 RPS5 RPS7A RPS7B RPS9B RRP9 RSP5 RVB1 RVB2 SPT5 SRM1 SSB1 SSZ1 SUI1 SUI2 SUI3 TIF11 TIF3 TIF34 TIF5 TMA20 YTM1 ZUO1 |

Third, the estimated half-lives agreed with literature-derived values that originate from targeted validation experiments (**Figure 3**, **Suppl. Table S1**, as taken from ref. (Christiano et al., 2014). The Christiano and Silva datasets also correlated weakly, but significantly with an orthogonal dataset on protein half-lives estimated by western blotting (Belle et al., 2006)(**Suppl. Figure S5**, p-value<0.01). This correlation is remarkable given that the western blot data was obtained with an invasive method using translation inhibition, a very short 45 min time course and only three measurement points.

**Figure 3:**
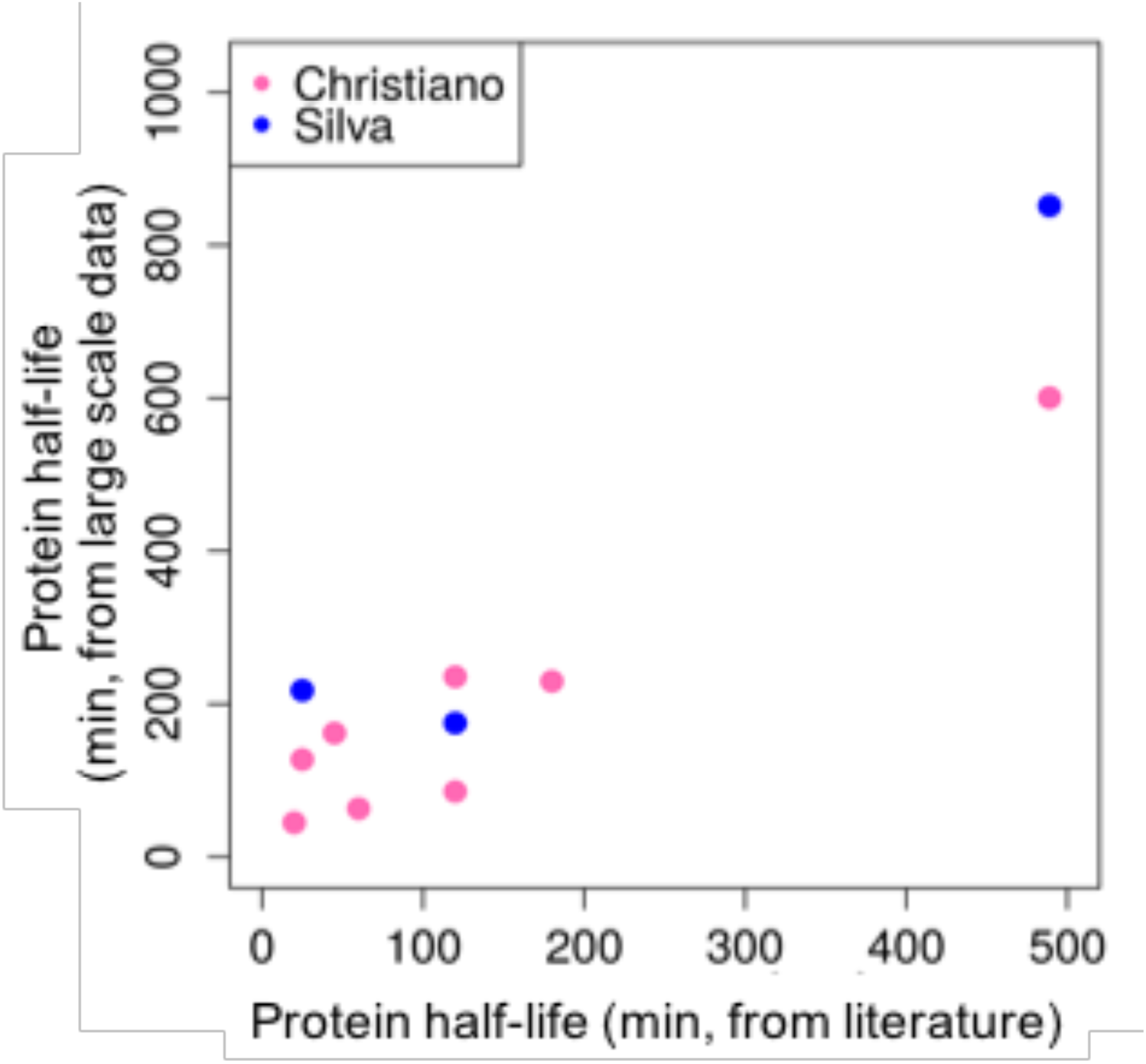
Validation of proteins with literature values. We compared the doubling time adjusted *T*_1/2_ to ten protein half-lives found in the 20 literature (**Suppl. Table S1**). The adjusted *T*_1/2_ values agree with the literature values, but cover a small range of values. Due to strict filtering, neither Martin-Perez dataset had data points amongst the literature confirmed values.

Fourth, as expected, protein half-lives correlate positively with abundance, both for binned and unbinned data (**Figure 4**). The correlation was weaker but still present for measures of codon adaptation which usually approximates abundances well, but not so for length which is known to inversely correlate with protein concentration (**Suppl. Figure S5**). Protein half-life estimates from the Christiano study showed a weak anti-correlation with both the presence of sequence degrons and intrinsically disordered regions: the more degrons or disordered regions a protein had, the less stable it appeared to be (**Suppl. Figure S5**). No searchable motif existed for ubiquitination (Kim et al., 2011)(*not shown*). We observed some correlations with amino acid frequencies (**Suppl. Figure S6**).

**Figure 4:**
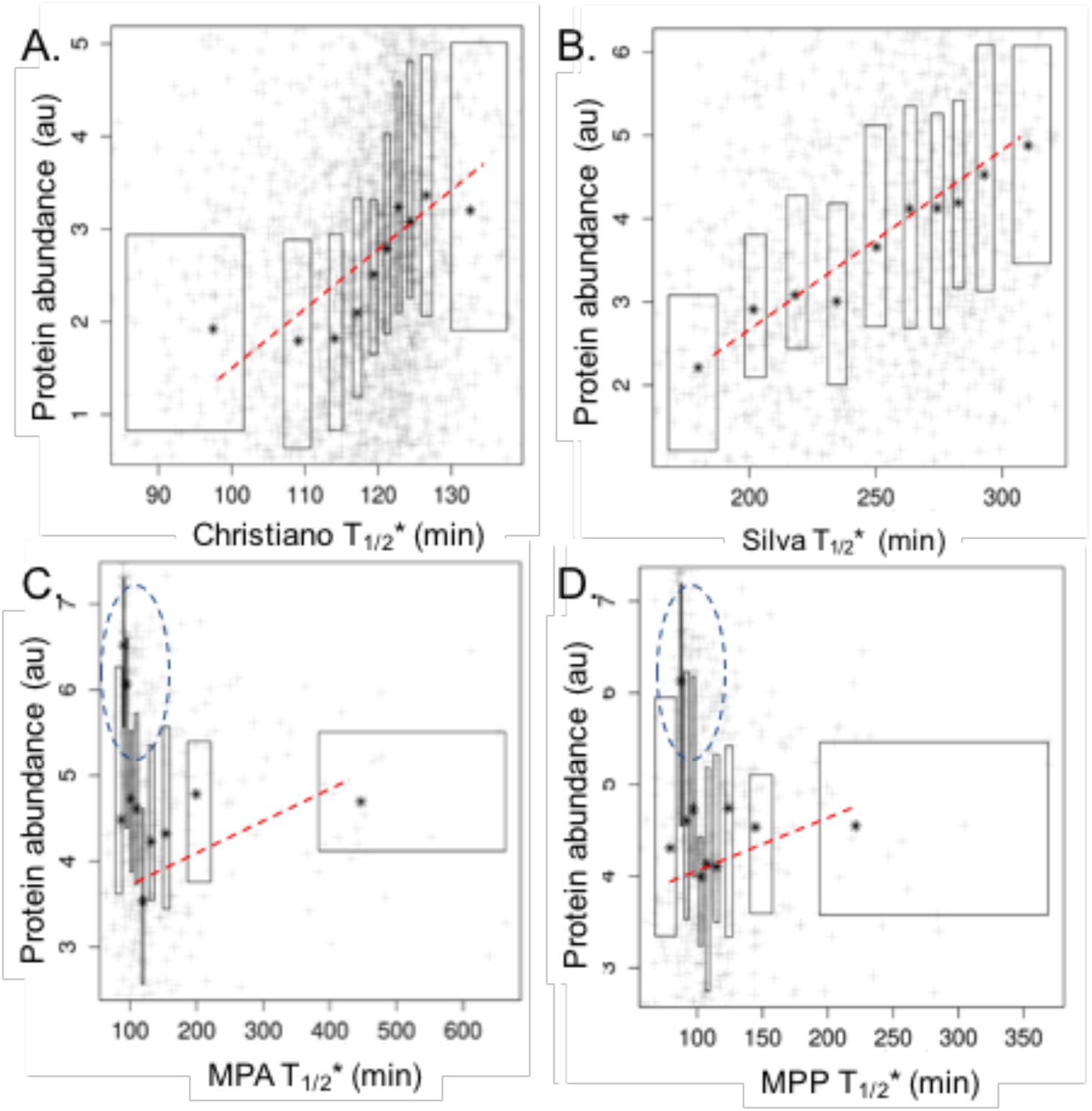
Protein half-lives correlate positively with abundance. The scatter plots show the the relationship between unadjusted protein half-lives (*T*_1/2_*) from the four datasets and protein abundance. Protein concentration data was obtained from ref. Csardi et al. (2015) and represents the corrected average over many datasets. Data were rank-ordered and then split into ten equal size bins. Black stars indicate the median values in each bin. Boxes indicate the second and third quartiles within the bins; stars indicate the median values. Correlations were calculated for all data points for Pearson’s r and Spearman’s ρ. Binned median values correlated significantly in all datasets with a p-value<0.01. Correlation coefficients for all values are: **A.** Christiano T_1/2_* vs. protein abundance; r = 0.21 (p< 0.01) and ρ = 0.33 (p < 0.01). **B.** Silva T_1/2_* vs. abundance; r = 0.39 (p < 0.01) and ρ = 0.39 (p < 0.01). **C.** Martin-Perez Prototroph (MPP) T_1/2_* vs. abundance; r = −0.03 (p = 0.56) and ρ = −0.22 (p < 0.01). **D.** Martin-Perez Auxotroph (MPA) T_1/2_* vs. abundance; r = 0.02 (p = 0.68) and ρ = −0.12 (p < 0.01). For the Martin-Perez datasets, a set of short-lived proteins (dotted oval) is enriched in ribosomal proteins. Extended results, e.g. on Codon Adaptation Index, are shown in **Suppl. Figure S5 and Suppl. Dataset 1.**

### Highly abundant proteins show inconsistent behavior

All four Martin-Perez datasets (duplicates for two experiments) displayed a set of short-lived proteins with varying abundance that were distinct from general trends in the data (**Figure 4**). This outlier set was surprising given the high quality of the data and its high reproducibility across replicates (Martin-Perez and Villen, 2015). Since this outlier set was enriched in highly abundant proteins (**Suppl. Figure S7**), and specifically ribosomal proteins, we examined half-lives of ribosomal proteins in the data in more detail.

In general, ribosomal proteins are of high-abundance and would therefore be expected to be long-lived compared to other proteins (**Suppl. Table S2**). Indeed, this expectation is met in the Christiano and Silva datasets on *S. cerevisiae* in this study, for *S. cerevisiae* and *Schizzosaccharomyces pombe* (Belle et al., 2006) (Christiano et al., 2014), for *Escherichia coli* (Schleif, 1967) and for mammalian cells (Schwanhausser et al., 2011). Surprisingly, the Martin-Perez data and for baker’s yeast examined in time series experiment using labeled nitrogen sources by Helbig et al. (Helbig et al., 2011) show a different picture: ribosomal proteins have comparatively short half-lives. To understand the reason behind this difference, we carefully compared the experimental setup for the Martin-Perez and Helbig studies to the other yeast experiments (**Suppl. Table S2**). There was no apparent reason for the differential behavior based on the amino acid dependence (proto-vs. auxotrophy), the culture type (batch vs. continuous culture), the prior labeling time, the type of label switch (heavy to light or *vice versa*), or the length of the overall time series experiment (**Suppl. Table S2**). The only weak difference that we detected lied in the exact method by which the medium was switched, i.e. *via* pelleting or inocculation.

However, we were able to demonstrate consistency of the stabilities of ribosomal proteins *amongst each other* (**Figure 5**). When we compared half-lives that were rank-ordered amongst ribosomal proteins, we found substantial agreement amongst the datasets (**Figure 5**). For example, the stalk proteins RPP0, RPP1A and RPP2A are particularly long-lived. Conversely, RPL10, RPL3, RPS7B, and the paralogs RPL7A and B are particularly short-lived compared to other proteins (**Figure 5**). The short half-live of these ribosome subunits might also link to their roles in late stages of ribosome biogenesis, consistent with the function enrichment that we observe (**Table 2**). Notably, the difference between the least and most stable ribosomal proteins in **Figure 5** was as small as eight minutes in unadjusted half-life (in the Martin-Perez datasets).

**Figure 5:**
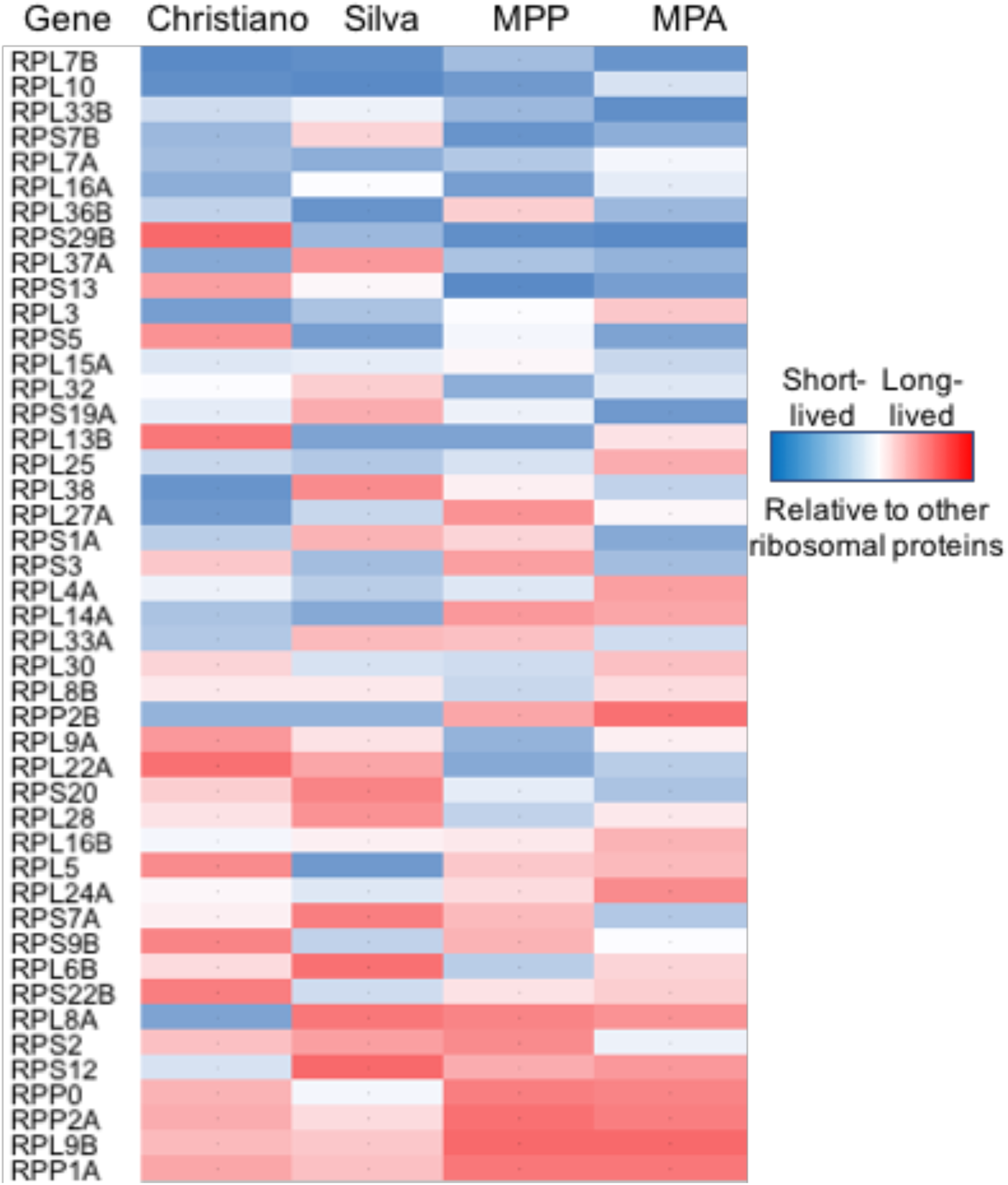
All experiments resolve differences in half-lives across ribosomal proteins. Ribosomal proteins are relatively rapidly turned over in the Martin-Perez datasets compared to non-ribosomal proteins (**Figure 4**). However, within ribosomal proteins the measurements of stabilities are consistent with other estimates. The figure shows only those ribosomal proteins with estimates in all four datasets. All protein half-lives are ranked within the shown set. The coloring denotes stability *relative* to other ribosomal proteins in the same set, it does not reflect absolute values. MPP – Martin-Perez Prototroph; MPA – Martin-Perez Auxotroph

## Conclusion

We consolidated very different half-life estimates for yeast proteins from identical conditions. We showed that, using consistent data post-processing and cutoffs, the datasets behave very similarly (**Suppl. Figure S3**) and common principles emerge. Degradation of many yeast proteins can be approximated by first-order decay, but true degradation functions might require more complex models (McShane et al., 2016). Most proteins are removed from the cell via dilution by cell division. After accounting for this dilution effect, i.e. differential doubling times, the median half-lives across the datasets were large, from 7.5 to 40 hours. Within a dataset, protein half-lives varied by an order of magnitude.

With these estimates, proteins are much more stable than mRNAs (Munchel et al., 2011; Neymotin et al., 2014; Sun et al., 2012). This high stability had been reported by Christiano et al. (Christiano et al., 2014), but not by earlier work suggested that used translation inhibitors (Belle et al., 2006) or recent work comparing prototroph and auxotroph strains (Martin-Perez and Villen, 2015). It implies that most of the decrease in pre-existing proteins arises from dilution by cell division and proteasomal degradation is slow. Interestingly though, even after accounting for cell doubling times, protein half-lives correlated with growth, suggesting that proteasomal degradation partially adjusted to growth rate (**Figure 1D**).

We observed consistent protein half-life characteristics across the datasets, correlation with literature-confirmed values, function enrichment, correlation with protein concentrations and some other sequence characteristics. The correlation with abundance and codon usage suggested that the cell efficiently produced higher steady-state concentrations of protein by an increase in both translation and protein stability. We observed an inverse relationship of half-life with the presence of destabilizing disordered regions and degrons (**Suppl. Figure S5**) – this relationship had been hidden in many previous studies.

We showed that these results are independent of the post-processing method, i.e. the method by which one accounts for sampling depth or technical variation between measured time points, and scaling of the data. Due to the remaining noise (**Suppl. Figure S8**), values for individual protein stabilities need to be considered with care. However, the methods were still able to discern relative differences in ribosomal half-lives as small as a few minutes.

These differences matched known observations and roles of ribosome subunits (Mathis et al., 2017). For example, the short half-life of RPL10 had first been reported in the 1970s, is conserved in mammals (Lastick and McConkey, 1976, and might be regulated by extensive ubiquitination {Bengsch, 2015 #8275; Mathis et al., 2017)(**Figure 5**). Ubiquitination was also observed for the short-lived RPS7 in **Figure 5** (Bengsch et al., 2015). Further, RPL10’s short half-life was consistent with its function: RPL10 controls the switch between the non-rotated and rotated state of the ribosome during elongation, and the protein has to be rapidly available and removed. In contrast, subunits of the stalk (RPP0, RPP1A, RPP2A, **Figure 5**) are not easily interchangeable and therefore expectedly long-lived. Finally, the two paralogs RPL33A and B display an interesting half-life difference: RPL33B is short-lived, while RPL33A is consistently long-lived across the four datasets. The rpl33a null mutant exhibits slow growth, the rpl33b mutant grows normally, rpl33a rpl33b double null mutant is inviable (Cherry et al., 1998; Hellerstedt et al., 2017; Wong, 2017), suggesting that the proteins might have overlapping, but slightly differing functions regulated by differential turnover.

In sum, several studies have examined protein half-lives in yeast and our meta-analysis has revealed common trends. The knowledge from these studies should be used to refine our view of protein stability, for example with respect subtle differences between protein half-lives. It should also be used to design future experiments that avoid discrepancies observed previously. Most importantly, half-life estimates need to distinguish clearly between estimates with or without the contribution of cell division. While accounting for cell doubling time does not change the ranking of half-lives across proteins, it has a non-linear and substantial effect on the half-life value (**Table 1**). Further, experiments need to cover two or more cell cycles, to avoid overestimation of values for stable proteins. Sampling four time points is sufficient for approximations of first-order decay which in turn provides good working estimates of half-lives. When monitoring half-lives over many hours, the non-exponential nature of the decay confounds analysis, and experiments will need denser sampling to model the true decay function. In other words, simple experiments can provide good working models of protein stability, but estimating exact half-lives, in particular of long-lived proteins, remains challenging and requires complex approaches.

## Methods

### Proteomics experiments

Quantitative pulsed Stable Isotope Labeling by Amino Acids in Cell Culture (SILAC) experiments were performed in cells grown at 30 C in synthetic dextrose medium containing amino acid dropout depleted in arginine and lysine (Sunrise Science). SILAC media were supplemented with light or heavy isotopes of arginine and lysine (L-Arg6 ^13^C; L-Lys8 ^13^C, ^15^N – Cambridge Isotopes). Yeast cells (GMS413) were grown in light medium for at least ten generations prior to transferring to heavy medium. Cells were then collected by centrifugation after 30, 60, 150 and 240 min. Cell disruption was performed by glass-bead agitation at 4 C in Urea buffer: 8 M Urea, 50 mM Tris-HCl pH 7.5, 150 mM NaCl, and 1x EMD Millipore protease inhibitor cocktail set I. The extract was cleared by centrifugation and protein concentration was determined by Bradford assay (BioRad). Protein preparation for proteomics analysis was performed as previously described (Silva et al., 2015). Next, 200 μg of tryptically digested proteins were fractionated at high pH (20 mM ammonium formate buffer, pH 10) reverse phase chromatography in a Kinetex 5μm EVO C18 100 Å column. Eluted peptides were non-linearly combined in 12 fractions and loaded for LC-MS/MS analysis. For each time point, 12 fractions were separated on an Agilent Zorbax 300 Stablebond-C18 column (3.5 μm, 0.075 x 150 mm) by reverse-phase chromatography for 160 min with a gradient of 5 to 60 % acetonitrile performed with an Eksigent NanoLC 2DPlus liquid chromatography system. The eluted peptides were in-line injected into an LTQ-Orbitrap Velos mass spectrometer (Thermo Scientific). Data-dependent analysis was performed at a resolution of 60,000 on the top 20 most intense ions from each MS full scan with dynamic exclusion set to 90 s if *m/z* acquisition was repeated within a 45 s interval. Mass Spectrometry (MS) MS1 data was acquired at the FTMS orbitrap mass analyzer with target ion value of 1e6 and maximum injection time of 500 ms. MS2 data was acquired at the ion trap mass analyzer with target ion value of 1e4, maximum injection time of 100 ms, isolation window of 2 m/z, and CID normalized collision energy of 35.

### Primary data analysis

The output RAW data was processed using MaxQuant suite (v. 1.3.0.5) to identify and quantify protein abundance (Tyanova et al., 2016). The spectra were matched against the yeast *Saccharomyces cerevisiae* database (Boutet et al., 2016; Consortium, 2015). Protein identification was performed using 20 ppm tolerance at the MS level (FT-mass analyzer) and 0.5 Da at the MS/MS level (Ion Trap analyzer), with a posterior global false discovery rate at 1% based on the reverse sequence of the yeast FASTA file. Up to two-missed trypsin cleavages were allowed, oxidation of methionine and N-terminal acetylation were searched as variable post-translational modification, and cysteine carbamidomethylation as fixed. The minimum number of SILAC peptide pairs used to quantify the protein abundance was set to two.

The dataset is labeled ‘Silva’ throughout this study. All raw and primary analysis files are freely available from the PRIDE database, identifier PXD005956. (For reviewing purposes: Username: Password: Q9cTf30N)

### Acquisition of published data

We obtained primary proteomics datasets, i.e. the MaxQuant output, from two published studies, labeled ‘Christiano’ and ‘Martin-Perez’ for the first authors, respectively (Christiano et al., 2014; Martin-Perez and Villen, 2015). Each study had conducted a SILAC proteomics experiment similar to those described above measuring time points as listed in **Table 1**. Martin-Perez et al. measured additional time points considered here as described in the **Results** section.

### Estimating cell doubling time

Cell doubling times *T*_*CC*_ for the Christiano and Martin-Perez studies were taken from the respective publications (**Table 1**)(Christiano et al., 2014; Martin-Perez and Villen, 2015). For the Silva dataset, we measured doubling times by monitoring the OD over time. As the *T*_*CC*_ varied slightly between time points, we scaled each time point to the measured *T*_*CC*_ between this time point and the previous time point.

### Secondary data processing

To enable fair comparisons across the datasets, we selected equal number of time points (four) and measurement periods (∼240min) across the studies (**Table 1**). To evaluate the robustness of the results, the calculations were repeated for extended studies. We obtained similar results for the MPP data after both truncating the time series at 120 min and extending the time series to 360 min (**Suppl. Figure S2**, **Suppl. Table S3**). We tested three post-processing methods to account for variation in mass spectrometry measurements across time points. All calculations were performed on an x86-64 GNU/Linux desktop using R version 3.3.2.

*Unscaled data.* For each protein, we performed a linear regression on *ln*(1+*P*_*new*_/*P*_*old*_) at each time point. This transformation corresponds to an exponential decay function.

*Ratio scaling.* To control for batch effects in which unexpectedly large or small intensities are observed for all proteins at one experimental time point, we scaled all protein intensities at each time point to the sum of all intensities at that time point. Because we expect a small number of highly abundant proteins to disproportionately contribute to the intensity sums, we compute scaling factors using only sums of the bottom 95% quantile. Additionally, we fit the intensity sums to an exponential decay function. For a SILAC protein degradation experiment, the total MS intensity is expected to stay constant, but the fraction of old proteins, *P*_*old*_, should decrease from 1 to 0 and the fraction of new proteins, *P*_*new*_, should increase from 0 to 1 over time, respectively. Thus instead of scaling intensities to a constant expected value, we scaled intensities to a linear model for *ln*(1+*P*_*new*_/*P*_*old*_).

*Metagene scaling.* In another scaling method, we constructed a metagene using the most stable 5% of proteins in each dataset. We then calculated the average *P*_*old*_ and *P*_*new*_ at each time point for these genes, then scaled all protein intensities in the dataset to the metagene values.

Ratio scaling returned the most significant genes for all datasets, and we used this method for all downstream analysis. The method used did not systematically affect the experimental outcome (**Suppl. Figure S1**).

### Modeling of protein half-lives

Pulsed SILAC is a proteomic technique which typically consists of growing auxotrophic cells in medium containing isotopically labeled amino acids, then switching cells to a medium containing amino acids with a new isotopic label at time (*T*_0_). Note that prototrophic cells were used in the Martin-Perez dataset (Martin-Perez and Villen, 2015). Since the use of heavy and light isotopes changed across the three studies, we chose to refer to peak intensities *P*_*old*_ and *P*_*new*_ when discussing the initial (‘old’) and pulsed (‘new’) isotope. Proteins containing the old label are assumed to have been synthesized before (*T*_0_), and proteins containing the new label are assumed to have been synthesized after (*T*_0_). Protein degradation can in theory be monitored simply as the disappearance of *P*_*old*_ without considering the SILAC ratios of *P*_*old*_/*P*_*new*_. However, the estimates using *P*_*old*_ alone are far less precise than those based on the ratio (*not shown*).

Protein half-life is commonly assumed to follow exponential decay. To establish which proteins can be well-modeled by exponential decay, we followed a procedure developed by Schwanhaeusser et al. and Christiano et al. (Christiano et al., 2014; Schwanhausser et al., 2011). The procedure is explained in the **Suppl. Notes**.

*Error estimates.* As established by Christiano et al. (Christiano et al., 2014), each protein’s fit to exponential decay can be modeled by the leave-one-out cross-validation (LOOCV) error and a general R^2^. LOOCV consists of recalculating the linear model of the half-life model with one data point left out. The generated model predicts a value for the omitted time point, then calculates the error between the predicted and actual value. This procedure iterates over each time point for the protein. The algorithm reports the mean error of iteration over all time points. The LOOCV was implemented using the cv.glm() function from the boot R package. R^2^ values were calculated from the linear regression using the summary.lm() function.

All data including error estimates are provided in **Supp. Dataset 1**. Both LOOCV and R^2^ depend on the number of data points considered which is the reason for us to choose four time points and similar measurement periods for each dataset to enable comparisons. **Suppl. Figure S3** shows the distributions of the errors after these corrections. Based on these distributions, we chose an LOOCV < 0.01 and R^2^ > 0.9 as common thresholds for all three datasets to consider a protein well-modeled by an exponential decay function.

Combining replicates. MPP and MPA both included two technical replicates. To obtain a consensus dataset for each, we filtered both replicates for significant proteins. For proteins that were significant in both replicates, T_1/2_ and T_1/2_* were obtained by taking the average value of both replicates.

### Analysis of protein properties

We performed Gene Set Enrichment Analysis (GSEA) on each dataset (Subramanian et al., 2005). GSEA determines whether a given gene set is overrepresented at the top of an ordered list of genes. To enrich for long-lived proteins, we order the proteins in a dataset by *T*_1/2_* with *T*_1/2_* decreasing. To enrich for short-lived proteins, we ordered the proteins in a dataset with *T*_1/2_* increasing. Estimates of absolute protein concentrations were taken from a publication by Csardi et al. (Csardi et al., 2015). Amino acid frequencies and codon adaptation indices were taken from the protein_properties file from SGD (Cherry et al., 1998; Hellerstedt et al., 2017; Wong, 2017). Scores for PEST sequences and intrinsically disordered regions were calculated using the EPESTFIND and DiEMBL tools, respectively, which are part of the EMBOSS package.

## Acknowledgements

CV is grateful to John C. Price for helpful discussions. CV acknowledges funding by the NIH (R01 GM113237), US National Science Foundation EAGER grant MCB-1355462, the NYU Whitehead Foundation, and the Zegar Family Foundation Fund for Genomics Research at New York University. This work was supported in part by the US National Institutes of Health K99 award ES025835 (GMS). The funders had no role in study design, data collection and analysis, decision to publish, or preparation of the manuscript.

## Supplementary files

*Supplementary Notes.*

Additional methods, figures, and tables.

## Supplementary Dataset 1

Protein half-lives for all datasets. Missing values indicate proteins that were absent from a dataset. t.half.star is T_1/2_* for the dataset. t.half.adjusted is T_1/2_ adjusted for T_CC_ for the dataset. is.significant is a logical value indicating whether the half-life calculation meets our significance cutoff of LOOCV error < 0.01 and R^2^ > 0.9. pearson.p.value is the p value given by the Pearson correlation of protein time points. FDR is pearson.p.value adjusted for false discovery rate.

## Supplementary Dataset 2

GSEA output for enriched GO Biological Process for each dataset.

## Supplementary Dataset 3

R scripts for calculations

